# Connecting the legs with a spring improves human running economy

**DOI:** 10.1101/474650

**Authors:** Cole S. Simpson, Cara G. Welker, Scott D. Uhlrich, Sean M. Sketch, Rachel W. Jackson, Scott L. Delp, Steve H. Collins, Jessica C. Selinger, Elliot W. Hawkes

## Abstract

Spring-like tissues attached to the swinging legs of animals are thought to improve running economy by simply reducing the effort of leg swing. Here we show that a spring, or ‘exotendon,’ connecting the legs of a human runner improves economy instead through a more complex mechanism that produces savings during both swing and stance. The spring increases the energy optimal stride frequency; when runners adopt this new gait pattern, savings occur in both phases of gait. Remarkably, the simple device improves running economy by 6.4 ± 2.8%, comparable to savings achieved by motorized assistive robotics that directly target the costlier stance phase of gait. Our results highlight the importance of considering both the dynamics of the body and the adaptive strategies of the user when designing systems that couple human and machine.

## Introduction

Running expends more energy than any other commonly used form of locomotion, including walking, swimming, and flying [1]–[3] (Fig. 1A). In running humans, only a small amount of the metabolic energy expended does net external work on the environment; this energy is used to overcome aerodynamic drag and represents less than 8% of the total energy expended (Fig. 1B) [4], [5]. The remaining energy is ‘wasted’ in the sense that it is expended by processes that do no useful external work on the environment. According to studies that attempt to partition the energy expended by these processes, most of the wasted energy (65-82%) is used to accelerate and brake the center of mass, both vertically and fore-aft, a process that occurs each stance phase [6]. A smaller portion is used to swing the legs [6]–[8], with the current best estimate at 7% [6].

**Fig. 1.**
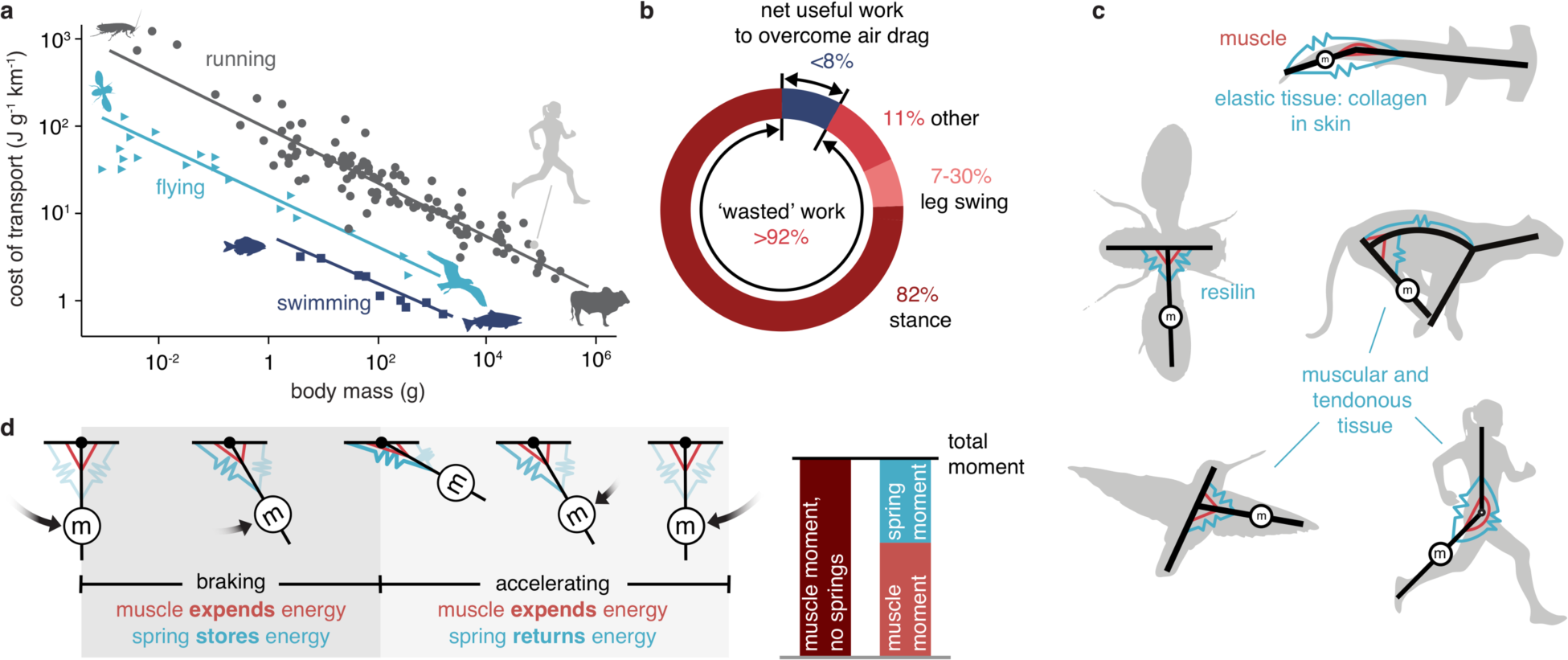
Energetics and mechanics in running animals. **a**, Cost of transport as a function of body mass [3], [30], [31] shows that running (grey circles) is less efficient than swimming (dark blue squares) and flying (light blue triangles). **b**, Only a small fraction of the energy expended in running does useful work on the environment to move against air resistance [4], [5]; the remainder is expended primarily to accelerate the center-of-mass, both vertically and fore-aft, during stance. Much less is used to swing the legs [6]–[8]. **c**, Elastic tissues are hypothesized to reduce the energy required to swing limbs. **d**, A pendular model of limb oscillation showing that a parallel spring (elastic tissue) can store energy during braking and return energy during acceleration, reducing required muscle moments.

Given the inefficiency of running, many devices have been designed to reduce a runner’s energy expenditure, with most targeting the costliest phase of gait—stance. These devices can be either active or passive. Active devices inject energy from an external source to reduce the amount of energy expended by the human, even while the total energy expended by the human-plus-device may increase. For example, exoskeleton robots use motors in parallel with human muscles [9], [10]. However, these active exoskeleton robots usually use offboard motors and power sources, which prevent them from being autonomous. Other examples of active devices include mechanisms with accelerated masses [11] and jet packs [12]. No actively powered device, however, has consistently reduced the energy required for a human to run while carrying the full weight of the device. Passive devices, in contrast, seek to reduce the energy required by the human to run by storing and returning energy to create a more efficient human-plus-device system. An early example is a running surface with stiffness tuned to minimize energy lost during impacts [13], [14]. Another passive assistance strategy involves using springs in parallel with the legs [15], [16], but this approach has yielded mixed results. Most recently, a shoe with carbon fiber springs embedded in the sole resulted in a 4% improvement in running economy [17], the largest savings for a self-contained system across all devices that target stance.

Notably, few devices have been designed that specifically target the energy expended for leg swing during running, even though numerous researchers hypothesize that passive elastic tissues in animals may reduce the energy required to oscillate limbs. Many quadrupeds have elastic tissues running along the top of the spine and front of the hip that are thought to assist spinal extension and hip flexion [18], [19] (Fig. 1C). Analogous passive elastic tissues are also in the skin of some fishes [20] and the wings of birds and insects [21], [22]. While no such mechanisms have been identified in humans, studies have correlated less flexibility in the legs and lower back with improved running economy [23]. This decreased flexibility might result from increased stiffness of passive elastic tissue spanning the relevant joints. For all of these examples, it is thought that the passive elastic tissues store and return energy during the oscillation of a limb, reducing the effort required to actively brake and accelerate the limb with muscles (Fig. 1D) [18] -[24]. Interestingly, recent work suggests that the savings resulting from assisting swing in humans are comparable to those seen when assisting stance [25], [26], despite the fact that the energy expenditure associated with stance is an order of magnitude larger [6]. Moreover, the savings associated with assisting swing may actually exceed the expected expenditure associated with swing [6], suggesting that the mechanism of savings when assisting swing is not understood.

## Results and Discussion

Here, we studied the effect of a simple spring connecting the legs of a running human on whole-body metabolic energy expenditure and found that running economy improved by 6.4 ± 2.8% (n=12, p=6.9×10^−6^, one-sample t-test), an amount comparable to devices targeting stance [9], [27]. We fabricated the externally mounted artificial elastic tendon, or ‘exotendon,’ from natural latex tubing (stiffness: 125 N/m, free length: 25% of the participant’s leg length, total mass: 22 g, see Materials and Methods) and attached each end of the device to the dorsal surface of the study participants’ shoes (Fig. 2, Supplemental Video). We measured metabolic cost using indirect calorimetry while twelve healthy adult recreational runners ran at 2.7 m/s on a treadmill. On each of two testing days, the participants completed two 10-minute runs with the exotendon, interspersed with two 10-minute natural runs without the exotendon for control purposes. We define a trial as a comparison between consecutive natural and exotendon runs, resulting in four trials over the two-day experiment. During the first trial of the first day, participants showed no metabolic savings when running with the exotendon compared to natural running. However, by the end of the second trial, participants were expending 3.8±5.4% less energy during exotendon running compared to natural running (n = 12, p = 0.034, one-sample t-test). Metabolic savings continued to increase on the second testing day, with all participants achieving savings by the end of the second trial of the second testing day (Fig. 3). To test for a possible placebo effect, we repeated the protocol with an exotendon whose stiffness was two orders of magnitude less than the original, thereby ensuring it could not provide substantial mechanical assistance. This placebo exotendon did not improve running economy (n=4, p=0.88, one-sample t-test, see Fig. S1). To test the safety and potential real-world applicability of the exotendon, four participants each ran 6 km on city streets (Fig. S2) with a modified exotendon (see Materials and Methods, Experiment 4); no tripping incidents occurred.

**Fig. 2.**
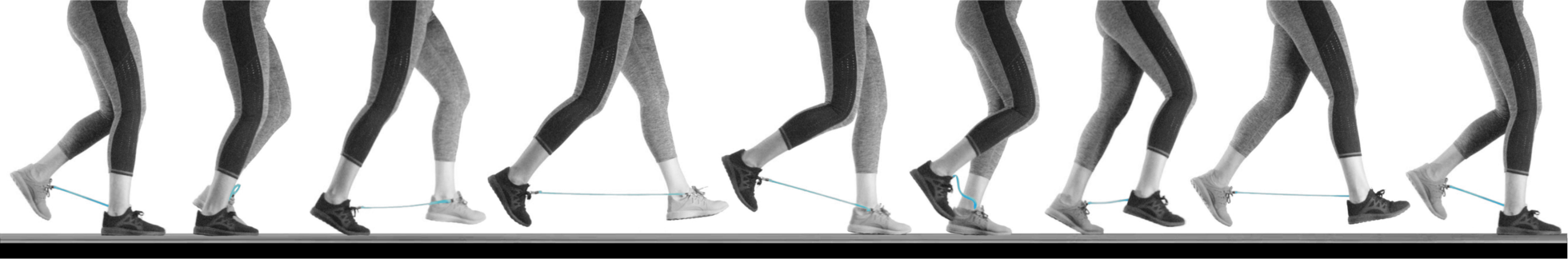
Time-lapse photographs of a runner using the exotendon. The length of the exotendon is tuned so that the device is long enough that it does not apply forces when the feet cross each other and does not break when the feet are far apart, yet short enough that it does not become entangled when the feet pass each other. Images span one complete gait cycle.

**Fig. 3.**
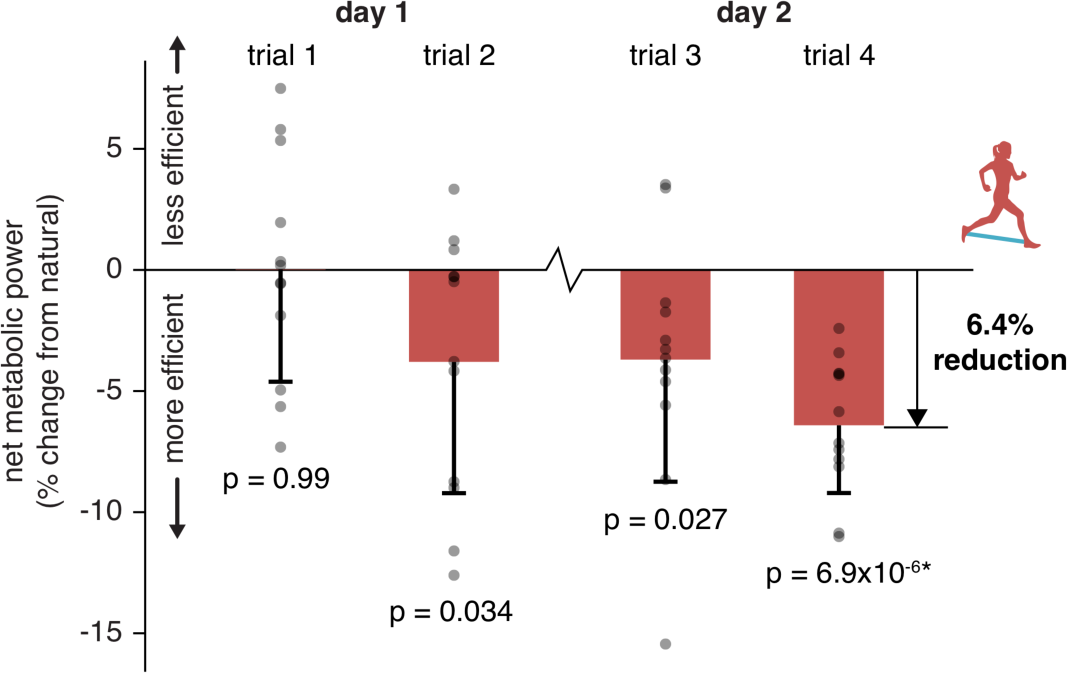
Reduced energy expenditure during exotendon running. On day 1, runners initially showed no change in energy expenditure (trial 1), yet showed reductions after running with the exotendon for 15-20 minutes (trial 2). Runners retained these savings across days (trial 3). After a total of 35-40 minutes of experience with the exotendon across both days, the greatest reductions in energy expenditure were evident (trial 4), with all runners (n=12) showing improved economy and average savings of 6.4 ± 2.8%. Error bars represent one standard deviation. Asterisks indicate statistical significance after Holm-Šidák corrections with confidence level α=0.05.

We hypothesized that our exotendon, designed to apply assistive moments primarily to the swing leg, also created savings during the stance phase of gait by increasing the energetically optimal swing rate and, in turn, reducing stance phase expenditures. This hypothesis can be explained as follows. During natural running, an increase in stride frequency tends to increase the energy expended to swing the legs [28] and decrease the energy expended during stance [29] (Fig. 4A). The stride frequency associated with minimum energy expenditure is thus a balance between these competing effects. However, if the exotendon reduces the energy expenditure associated with swinging the legs, the optimal stride frequency should increase. If runners adopt this higher stride frequency, expenditures associated with stance should decrease. In support of this hypothesis, we found that runners using an exotendon self-selected a higher stride frequency than when running naturally (8.9 ± 3.7%, n=12, p = 2.4*×*10^−4^, paired t-test, Fig. S3).

**Fig. 4.**
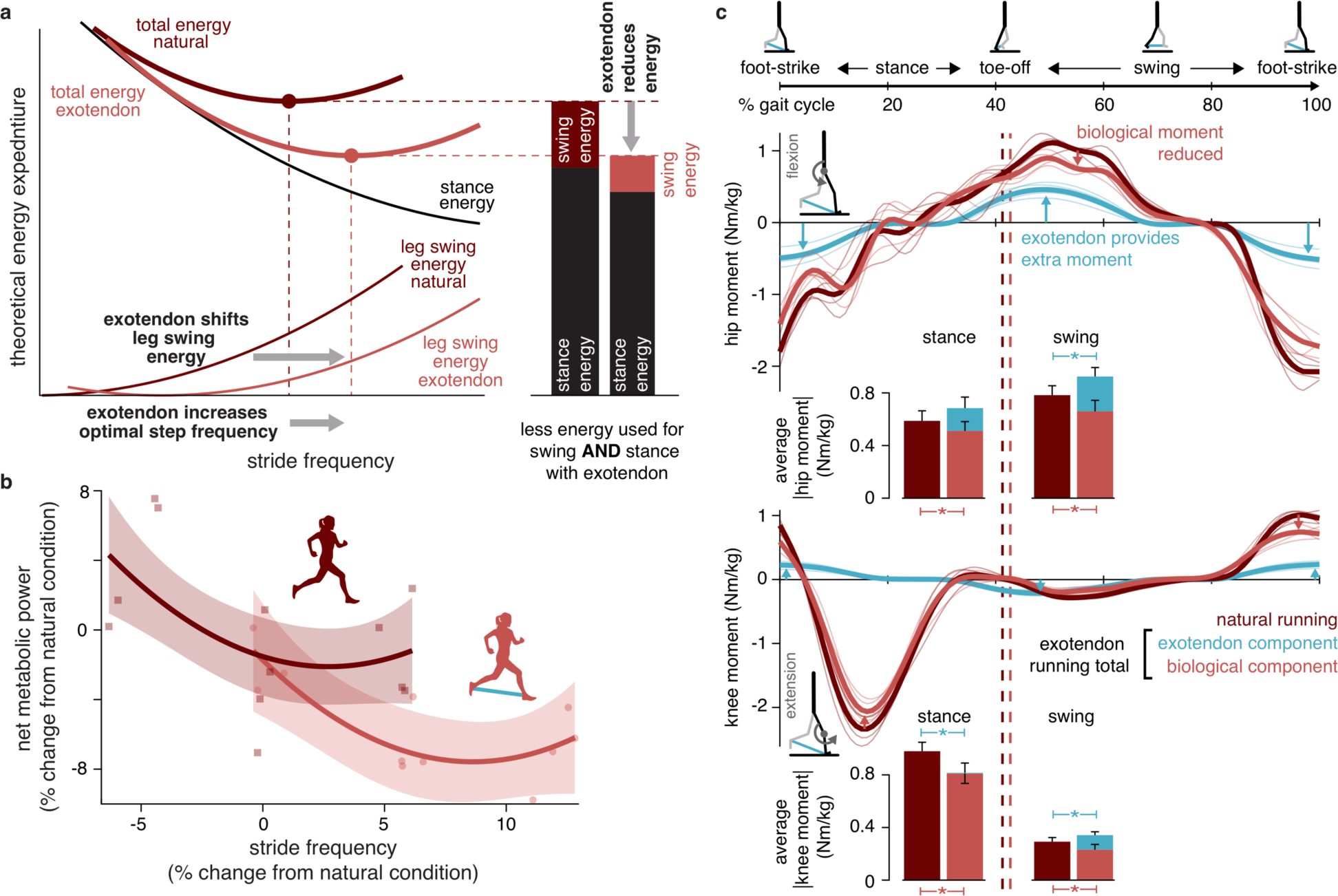
Exotendon mechanism of savings. **a**, Theoretically, runners choose an energetically optimal stride frequency (dark-red circle), which may result from a combination of processes that require more energy with increasing stride frequency, such as leg swing (dark-red thin line), and those that require less energy with increasing stride frequency, such as center-of-mass acceleration during stance (black thin line). We hypothesized that the exotendon shifts the leg swing curve (light-red thin line), increases the optimal stride frequency, and reduces total energy expenditure (including stance expenditure). **b**, In experiments, the exotendon increased the energetically optimal stride frequency (8.1%, p=3.7*×*10^−3^, paired t-test, n=4). Faded regions show the 95% confidence interval of curve fits. **c**, Biological moments during swing were reduced, likely due to the assistance of the exotendon, and biological moments during stance were reduced, possibly due to the increased stride frequency. Note that the exotendon can apply moments to the stance leg through the swinging leg.

To further test this hypothesis, we conducted additional experiments and found that the exotendon increases not only the self-selected stride frequency but also the energetically optimal stride frequency; it also reduces lower-limb joint moments and powers during both swing and stance. A subset of our participants (4 of 12) ran at a range of enforced stride frequencies both with and without the exotendon. We recorded kinematics and ground reaction forces to compute lower-limb inverse dynamics (Fig. 4C), as well as electromyography to capture lower-limb muscle activities (see Methods, Experiment 3). The exotendon significantly increased the optimal stride frequency (+8.1%, p=3.7×10^−3^, n=4, paired t-test), and all participants adapted toward the new optimum (Fig. 4B). The exotendon reduced biological hip and knee moments during swing (p=2.3×10^−3^, 2.5×10^−3^, respectively, paired t-test) and stance (p=4.1×10^−3^, 8.1×10^−5^, respectively, paired t-test) (Fig. 4C, Figs. S4 and S5). Interestingly, total knee moments during stance decreased (p=4.4×10^−4^, paired t-test), even when the exotendon was applying negligible moments, likely because runners adopted a higher stride frequency, as our hypothesis suggests. Corresponding reductions in muscle activities were not significant, possibly due to the low signal-to-noise ratio (Figs. S6 and S7). Overall, these results support our hypothesis that exotendon savings come from both swing and stance and highlight the interconnectedness of mechanics, energetics, and human behavior.

Assisting low-expenditure components of gait, such as leg swing, can result in greater-than-expected energy savings (see Fig. S8). During natural running, low-expenditure components can have high expenditures when a gait parameter, such as stride frequency, is changed outside of the preferred range. These sharp increases in expenditure act as a constraint, preventing adjustment to the gait parameter. During exotendon running, this constraint is relaxed, freeing the runner to reduce expenditures associated with the high-expenditure components of gait (such as stance) and achieve more efficient gait patterns overall. Critically, the associated savings can be large— much larger than would be expected from savings directly associated with a low-expenditure component of gait.

Our study shows how a spring designed to assist leg swing can significantly improve human running economy through a complex mechanism of savings. The device changes the relationship between stride frequency and energy expenditure, driving the runner to discover new locomotor strategies. This change in turn reduces effort during both swing and stance and results in overall greater efficiency than anticipated. Our exotendon could serve as an affordable and low-tech assistive device to improve human running performance, or a simple and robust intervention to further explore the complexities of human gait and human-machine interactions. More broadly, our study shows that a simple device can create unexpected and complex interactions between the dynamics of the body and the adaptive strategies of the individual—an important reminder for all who seek to augment humans.

## Acknowledgments

Thanks to Rachel Troutman for assistance running pilot studies, Michael Raitor for comments, insights, and enthusiasm, and Allison Okamura for always supporting her students, even if they stray far from her area of expertise.

## Funding

Funding for this work was provided by National Science Foundation Graduate Research Fellowships (CSS and SDU, grant #DGE-114747; CGW, grant #DGE-1656518), a Stanford Bioengineering Fellowship (CGW), a Stanford Graduate Fellowship (SDU), a Pfeiffer Research Foundation Stanford Interdisciplinary Graduate Fellowship affiliated with the Wu Tsai Neurosciences Institute (SMS), National Institutes of Health Big Data to Knowledge Center (SLD, grant #U54 EB0202405), the National Center for Simulation in Rehabilitation Research (SLD, grant #NIH P2C HD065690), and National Science Foundation (SHC, grant #CMMI-1818602).

## Author contributions

CSS and EWH conceived of the device. CSS, CGW, JCS, and EWH conceptualized the study. All authors contributed to experimental design. CSS, CGW, and SDU conducted the experiments. CSS, CGW, SDU, and SMS analyzed the results, with supervision from RWJ, SHC, JCS, and EWH. CSS, CGW, SDU, JCS, and EWH drafted the manuscript. All authors reviewed and edited the manuscript.

## Competing interests

The authors declare no competing interests.

## Data and materials availability

The data that support the findings of this study are available from the authors on request. Prior to final submission, we will also make data and musculoskeletal simulations available on SimTK.

## Methods

### Device design

We constructed our exotendons out of natural latex rubber surgical tubing (hollow cylindrical tubing, 0.95 cm outer diameter, 0.64 cm inner diameter). Each exotendon consists of a single length of tubing with a 1 cm loop at each end for attachment purposes. To make each loop, we folded the tubing, stretched the loop by hand, and wrapped the looped tubing tightly with electrical tape. Once released, the forces provided by Poisson expansion of the tubing supplement the adhesive, forming a secure connection. We then attached each loop to a 1.6 by 0.5 cm s-shaped stainless-steel carabiner that was clipped to the shoelaces of each participant. The length of the exotendon from end to end of each attachment loop was set to 25% of the participant’s leg length, measured as the distance from the top of the anterior superior iliac spine to the medial malleolus of the ankle. Through pilot testing, we found that this length was long enough to avoid breaking and short enough to avoid tripping during running. A completed exotendon is shown in Fig. 2 and the supplementary video.

Designing devices that reliably and accurately apply forces to the human body is a challenge. It often requires overcoming a myriad of difficulties including: aligning device and joint axes *(32)*, limiting added mass and materials to the body, and comfortably transferring force from rigid devices to often soft and deforming body segments (*33*). While we could have designed our device to attach more proximally, at the knee or hip for example, we found in piloting that the aforementioned challenges could be largely avoided by affixing our device to the shoes. In addition, attaching more distally on the leg offered two further advantages. First, due to a longer moment arm, the forces necessary to provide assistive moments to the limb were smaller than if the attachment points were more proximal. Second, more distal placements ensure the line-of-action of the spring is predominantly along the flexion-extension axis of the hip, minimizing adduction moments on the leg.

### Device design

_A total of 19 healthy young adults, with no known musculoskeletal or cardiopulmonary impairments, participated in the study (8 females; age: 24.9±2.7 years; height: 174.4±6.9 cm; weight: 67.3 ± 11.0 kg). The study was approved by the Standard University ethics board and all participants provided written informed consent prior to testing.

### Experimental protocols

We conducted four separate experiments to: determine if the exotendon improves running economy (Experiment 1), test for the possibility of a placebo effect (Experiment 2), determine the mechanism of energy savings (Experiment 3), and test the safety of the device during over ground running (Experiment 4).

#### Experiment 1 – Running economy

To determine if the exotendon improves running economy, we conducted an experiment to compare metabolic energy expenditure with and without the exotendon. Twelve participants (5 females; age: 24.7±2.9 years; height: 177.0±6.7 cm; weight: 69.3±11.4 kg) completed a two-day running protocol. On the first day of testing, we measured participants’ leg lengths and constructed personalized exotendons (25% of leg length). Participants were told that the exotendon was designed to improve running efficiency and were told to ‘relax into running with the device’ and try to ‘think about something else’ while running. The exotendons were then attached to participants’ shoes to allow for habituation to the device. This included a minimum of four 15 m over-ground walking and running trials and continued until participants verbally confirmed they were comfortable walking and running with the exotendon. Participants were then instrumented with indirect calorimetry equipment (Quark CPET, Cosmed, Rome, Italy) and completed a 5-minute quiet standing trial, during which baseline metabolic energy expenditure was measured. Participants then completed four 10-minute runs, with 5-minute rests between each, on a treadmill (Woodway, Waukesha, WI) at 2.67 m/s (10 minutes/mile). The runs alternated between ‘natural running’ (without the exotendon) and ‘exotendon running’ (with the exotendon), with the first running condition randomly assigned. The second day of testing was identical to the first for each participant. We define a trial as the comparison between consecutive natural and exotendon runs resulting in four trials over the two-day experiment. During runs, we recorded sagittal plane video, which we later used to determine runners stride frequency. However, in 5 of 12 participants, due to equipment availability, we recorded stride frequency using an accelerometer (Trigno IM, Delsys Inc., Natick, MA, USA) mounted on the dorsal surface of the foot.

#### Experiment 2 – Placebo effect

To determine if a placebo effect could explain the changes in running economy observed in Experiment 1, we conducted a separate experiment to compare metabolic energy expenditure during running with and without a placebo exotendon. Four naïve participants (2 females; age: 24±2.2 years; height: 168.3±2.5 cm; weight: 60.1±10.8 kg) completed a two-day running protocol that was identical to Experiment 1. The only difference was that participants ran with an exotendon that had a stiffness two orders of magnitude lower than the original exotendon (5 N/m vs 120 N/m, respectively), and therefore provided negligible assistive moments to the limbs. The length of the placebo exotendon was still set to 25% of participant leg length.

#### Experiment 3 - Mechanism

To determine how the exotendon reduces energy expenditure we conducted an experiment to test how running mechanics and muscle activity change during exotendon running. Four participants (2 females; age: 25.0±1.6 years; height: 179.5 ±7.4 cm; weight: 75.3±13.7 kg), randomly selected from the 12 that participated in Experiment 1, completed an additional third day of testing. During this testing day, kinematic, kinetic, electromyographical (EMG), and metabolic data were recorded during running with and without the exotendon at a range of stride frequencies.

All runs were completed at 2.67 m/s on an instrumented treadmill (Bertec Corporation, Columbus, OH, USA) to allow for collection of ground reaction forces (2000 Hz). Kinematic data were recorded at 100 Hz using a 9-camera optical motion tracking system (Motion Analysis Corporation, Santa Rosa, CA, USA). Anatomical reflective markers were placed bilaterally on the 2nd and 5th metatarsal heads, calcanei, malleioli, femoral epicondyles, anterior and posterior superior iliac spines, and acromion processes, as well as on the C7 vertebrae. An additional 16 tracking markers, arranged in clusters, were placed on the shanks and thighs of both legs. Markers on the medial malleoli and femoral epicondyles were removed following the static trial. EMG data were recorded (Trigno IM, Delsys Inc., Natick, MA, USA) at 2000 Hz from the following 15 muscles of a single limb: peroneus, soleus, medial and lateral gastrocnemii, tibialis anterior, medial and lateral hamstrings, gluteus medius and maximus, vastus lateralis and medialis, rectus femoris, sartorius, adductor group, and iliopsoas. EMG electrodes were placed in accordance to SENIAM guidelines (*34*). Metabolic power was measured using indirect calorimetry (Quark CPET, Cosmed, Rome, Italy).

To warm up, participants ran without the exotendon for 5 minutes on the treadmill. Next, participants completed a series of maximum voluntary contractions (MVCs) for later normalization of EMG signals. These MVCs included five maximum height jumps and five sprints (*35*), in addition to one isometric and three isokinetic maximum contractions of the hamstrings, adductor group, tibialis anterior, peroneus, hip flexors (with both a flexed knee and extended knee), and hip abductors. We then recorded motion capture marker positions and ground reaction forces during a static standing trial for later scaling of a musculoskeletal model. As in Experiment 1, participants were habituated to the device through a series of over-ground walking and running trials. Participants were then instrumented with indirect calorimetry equipment and a 5-minute quiet standing trial was recorded to capture baseline metabolic energy expenditure.

Participants then completed two 7-minute runs, one ‘natural running’ (without the exotendon) and one ‘exotendon running’ (with the exotendon), with a 5-minute rest between and the order randomly assigned. Kinematic, kinetic, EMG, and metabolic data were recorded. Self-selected stride frequency was computed during the last minute of each run from the instrumented treadmill force signals with a custom Matlab (Mathworks Inc., Natick MA) script. We will refer to these self-selected stride frequencies as *natural self-selected stride frequency* and *exotendon self-selected stride frequency*.

To determine how the relationship between stride frequency and metabolic power changed when running with the exotendon, participants next completed six additional 7-minute runs during which step frequency was prescribed using a metronome. Participants ran at three prescribed stride frequencies during both natural and exotendon running. For the natural running conditions, the following three stride frequencies were prescribed: i. the participant’s natural self-selected stride frequency; ii. the participant’s exotendon self-selected stride frequency, which was higher than the natural self-selected stride frequency; and iii. a stride frequency lower than the natural self-selected stride frequency. The change from the natural self-selected stride frequency to the lower stride frequency was set to the percent difference between the natural self-selected stride frequency and the exotendon self-selected stride frequency. For the exotendon running conditions, the following three stride frequencies were prescribed: i. the participant’s exotendon self-selected stride frequency; ii. the participant’s natural self-selected stride frequency; and iii. a stride frequency higher than the exotendon self-selected stride frequency. The change from exotendon self-selected stride frequency to the higher stride frequency was similarly set to the percent difference between the natural self-selected stride frequency and exotendon self-selected stride frequency.

#### Experiment 4 - Overground test

To test whether the exotendon can safely be used in the real-world, we conducted an experiment to monitor fall risk during outdoor running. Four participants (2 females; age: 27.8±1.3 years; height: 172.6 ±6.1 cm; weight: 66.7±8.0 kg), who had previous experience with the exotendon through pilot testing or participation in Experiment 1, ran with a modified exotendon for 6 km on suburban streets along with the experimenter. The route followed by the participants is shown in Fig. S1. The modified exotendon, which attached directly to the ankle via a compression brace, was reported to be more comfortable than the original exotendon, which attached directly to the shoelaces. Moving the attachment point off the shoes also reduced wear of the shoelaces caused by sliding of the carabiner. This small change in attachment point was not expected to have a significant effect on running economy. The number of tripping and falling incidents were recorded.

#### A Note on Device Optimization

In supplementary pilot experiments (not presented here), we did attempt to perform human-in-the-loop optimization to determine the optimal exotendon length and stiffness. Four participants (two who had previously completed Experiment 1 and two naïve participants) completed a protocol similar to that described in Zhang et al. (*10*) We were unable to identify length and stiffness combinations with better performance than our standard device in the four pilot participants. One possible explanation is that our chosen device parameters where indeed near optimal. Another possible explanation is that participants were more risk averse in running with our device and thus adopted control strategies that prioritized stability (not falling) over efficiency. Parsing the different effects of human-in-the-loop optimization is left for future work.

## Data analysis

### Experiment 1 - Running economy

#### Metabolics

We computed the gross metabolic power (energy expenditure) from indirect calorimetry *(36)* by averaging data from the last two minutes of each experimental run. Baseline metabolic power, calculated as the average metabolic power during the last two minutes of the rested standing trial, was subtracted from our gross metabolic power measures to get net metabolic power during each run. We then computed the percent change in net metabolic power, from natural running to exotendon running for each of the four trials. We then used two-tailed, one-sample t-tests, with a Holm-Šidák correction, to determine if percent changes in net metabolic power were significant. The results of these analyses are presented in Fig. 3.

#### Stride frequency

We manually determined average stride frequency from video recordings by counting the strides taken and dividing by the time required to take them. When foot mounted accelerometers were instead used to compute stride frequency, we bandpass filtered accelerometer data (4^th^ order, zero-phase shift Butterworth, 2-20 Hz), summed the X, Y and Z accelerations, identified peak accelerations, and computed stride frequency as one over the average time between peaks. Average stride frequency measures were not statistically different between the two measurement methods. The results of these analyses are presented in Fig. S3.

### Experiment 2 - Placebo effect

#### Metabolics

We performed the same metabolic analyses as described in Experiment 1. The results of these analyses are presented in Fig. S1.

### Experiment 3 - Mechanism

#### Metabolics

We performed the same metabolic analyses as described in Experiment 1 to determine the average net metabolic power for each run. To determine the effect of altering stride frequency, we computed the percent change in average net metabolic power for each enforced stride frequency, both with and without the exotendon, relative to natural running (without the exotendon and with no enforced stride frequency). For each participant, we then used least squares regression to find the best-fit quadratic curves relating net metabolic power to stride frequency for both exotendon and natural running. We then calculated the stride frequencies at the minima of the natural running and exotendon running best-fit curves, which we will refer to as the *natural optimal stride frequency* and the *exotendon optimal stride frequency*, respectively. To determine if the exotendon shifted the optimal stride frequency, we performed two-tailed paired t-tests comparing exotendon optimal stride frequency to natural optimal stride frequency. Using all participant data, we also solved for best-fit quadratic curves relating net metabolic power to stride frequency for both exotendon and natural running, and calculated the 95% bootstrap confidence intervals for these across participant curves. The results of this analysis are presented in Fig. 4b.

#### Musculoskeletal modeling

Joint-level kinematics, kinetics, and mechanical powers were computed using a modified musculoskeletal model (*37*) in OpenSim 3.3 (*38*). Of the original 37 model degrees of freedom, we locked 18 including ankle eversion, toe flexion, and all those associated with the arms, leaving us with a 19 degree-of-freedom model. We generated subject-specific models by scaling the generic model to match the anthropometry of each subject during a standing static trial. For scaling, ankle and knee joint centers were calculated as the midpoint of the calcanei markers and femoral epicondyle markers, respectively, while the hip joint centers were calculated using a regression model based on the marker positions of the posterior and anterior superior iliac spines *(39)*. After low-pass filtering the marker positions at 15 Hz (4th order, zero-phase shift Butterworth), we computed joint angles using the OpenSim inverse kinematics tool. This tool uses a weighted least squares algorithm to pose the model in a way that minimizes the error between model and experimental marker locations. Joint moments were computed using the OpenSim inverse dynamics tool, which uses ground reaction forces and moments, joint angles from inverse kinematics, and classical equations of motion to solve for intersegmental moments. The joint angles used as input were low-pass filtered at 15 Hz (6th order, zero-phase shift Butterworth), and ground reaction forces and moments were low-pass filtered at 15 Hz (4th order, zero-phase shift Butterworth).

We modeled the exotendon in OpenSim as a linear path spring with a deadband range equal to its slack length. The spring forces were applied to the calcaneous body of each foot at the location of the band attachment marker from the static trial. The length and stiffness of the modeled exotendon was scaled for each participant. Inverse dynamics were first computed without the modeled exotendon to determine the total joint moments required to produce the resultant motion and ground reaction forces, referred to as *exotendon running total moments*. Inverse dynamics were then recomputed with the modeled exotendon for all exotendon runs to determine the moments produced solely by biological muscle and tissue, referred to as the *biological moments*. The moments applied by the exotendon were computed as the difference between the exotendon running total moments and biological moments, referred to as *exotendon moments*. Powers were then computed at each joint by multiplying moments by angular velocities.

Participants’ average joint angles and moment as a function of gait cycle for the hip, knee and ankle were calculated from the last minute of each run. To do this, we averaged across strides after normalizing each stride time to 100% gait cycle, computed as the time from heel strike to subsequent heel strike on a single leg. Strides were excluded from these average trajectories if the value of the measure exceeded 5 standard deviations from the mean at any time point in the stride. This resulted in the removal of 3% of strides on average for all runs and participants. Joint powers were then computed from the averaged joint angles and moments for each participant. Joint powers and moments were then normalized to body weight and across-participant average trajectories were computed. These results are displayed in Fig. 4c and Fig. S4.

We next computed the average absolute natural running and exotendon running moments and powers (both biological and exotendon) during the stance and swing phases of gait. We tested for differences between natural running and exotendon running (again, both biological and exotendon) using two-tailed paired t-tests with Holm-Šidák corrections. The results of this analysis are presented in Fig. 4c and Fig. S5.

We performed these analyses, comparing joint moment and powers during the swing and stance phases of gait both with and without the exotendon, as a means of estimating effort during each phase. Metabolic power, measured using indirect calorimetry, is our most direct measure of energy expenditure, but cannot be used to distinguish stance and swing expenditures; their effects are intermingled during the long sampling period of indirect calorimetry. Instead we analyzed the mechanical requirements of the body (joint moments) during each phase of gait. Previous studies have shown strong correlations between metabolic power and joint moment *(40)*, but we note that reduced joint moments do not guarantee reduced metabolic rate *(41)*.

*Electromyography*. Electromyograms from each muscle were bandpass filtered at 30-500Hz (4^th^ order, zero-phase shift Butterworth), rectified, and then low-pass filtered at 6Hz (4^th^ order, zero-phase shift Butterworth) to create linear envelopes. Envelopes were then normalized to the peak signal from the MVCs (*35*) to compute muscle activities. We then averaged muscle activities across strides from the final minute of each run, then normalized to 100% of the gait cycle. Strides in which the muscle activities exceeded 5 standard deviations from the mean for any time point were excluded from the average curve. All remaining EMG signals were visually examined and excluded if they appeared corrupted. Overall, 8% were excluded, with no bias towards exotendon or natural running. All processing was performed using custom Matlab scripts. The results of this analysis are presented in Fig. S6. We also computed average muscle activities during the stance and swing phase of gait, both for natural and exotendon running. We used two-tailed paired t-tests with Holm-Šidák corrections to compare activity during natural and exotendon running, during both stance and swing. The results of these analyses are shown in Fig. S7.

---

**Fig. S1.**
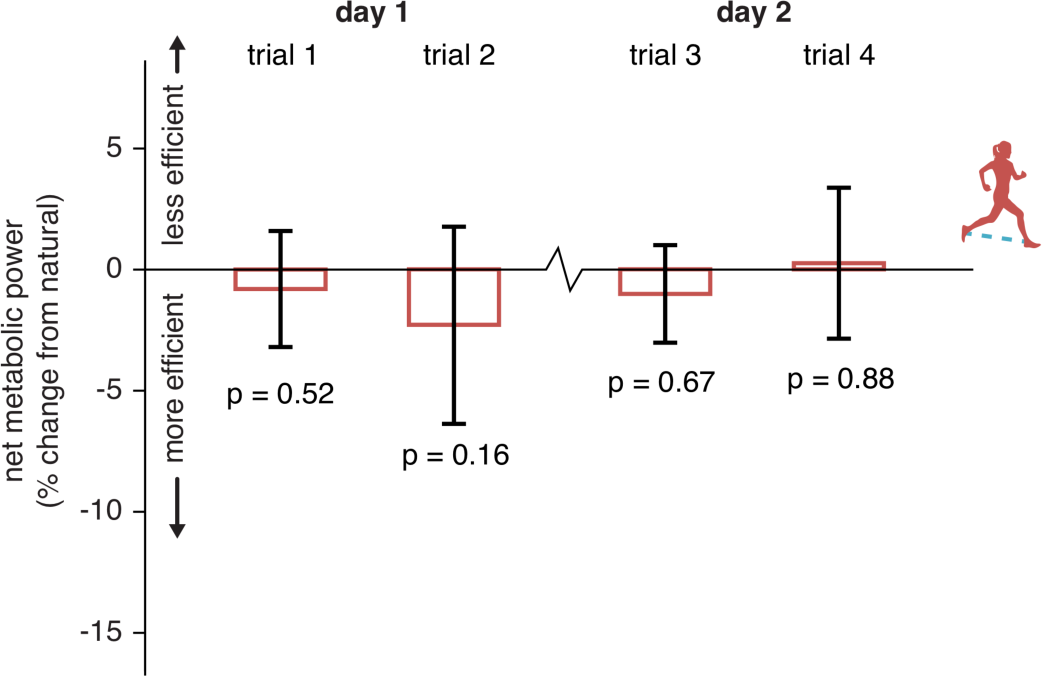
Placebo test results. Four participants completed the same protocol as the main experimental group, but were given an exotendon with stiffness less than 5% that of a normal exotendon. These participants showed no change in running economy, relative to natural running, with these placebo exotendons (two-tailed one-sample t-tests). Error bars represent one standard deviation.

**Fig. S2.**
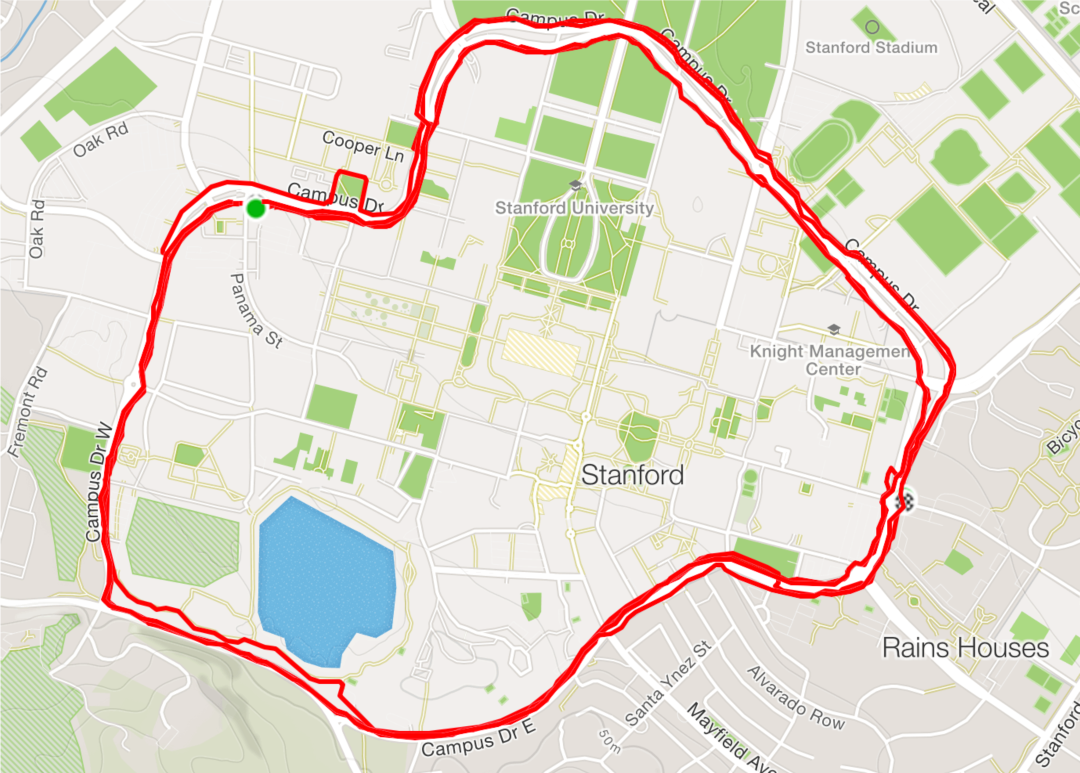
Overground running route. Four participants ran 6 km on city streets in modified exotendons to test the safety of the device during real-world over ground running.

**Fig. S3.**
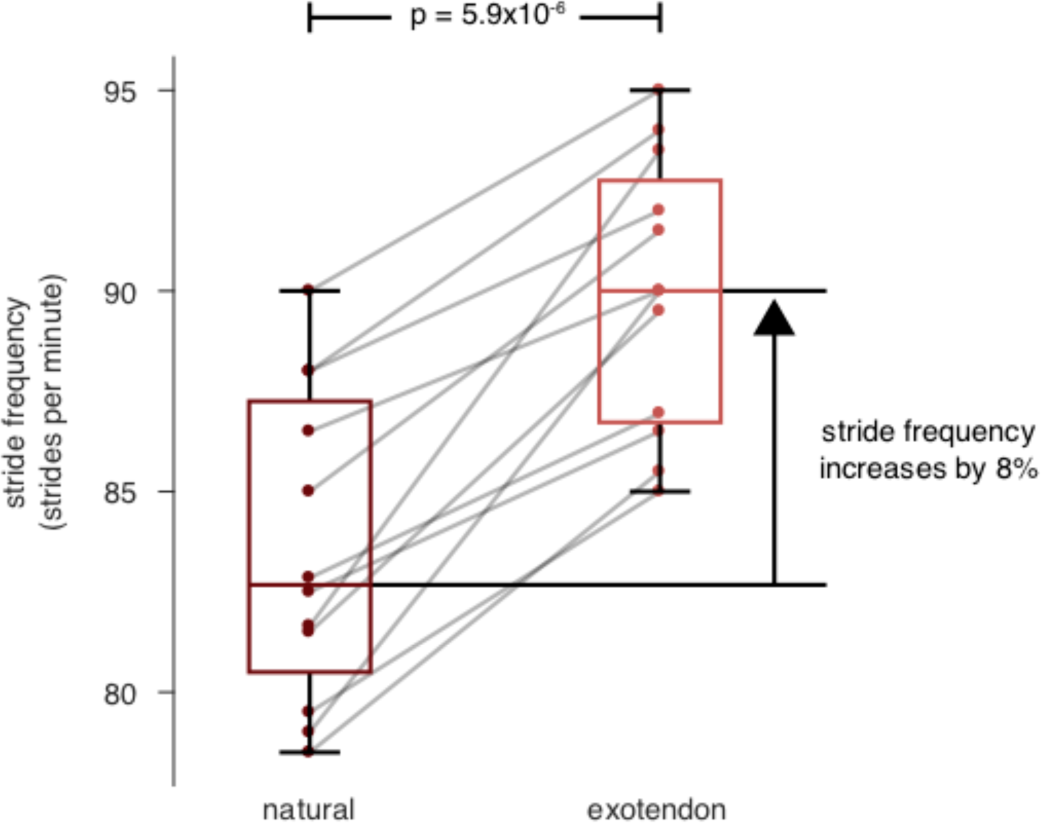
Stride frequency change. Participants took shorter, faster strides with the exotendon, increasing stride frequency by an average of 8% above that measured during natural running (p=5.9×10^−6^ two-tailed paired t-test, n=12).

**Fig. S4.**
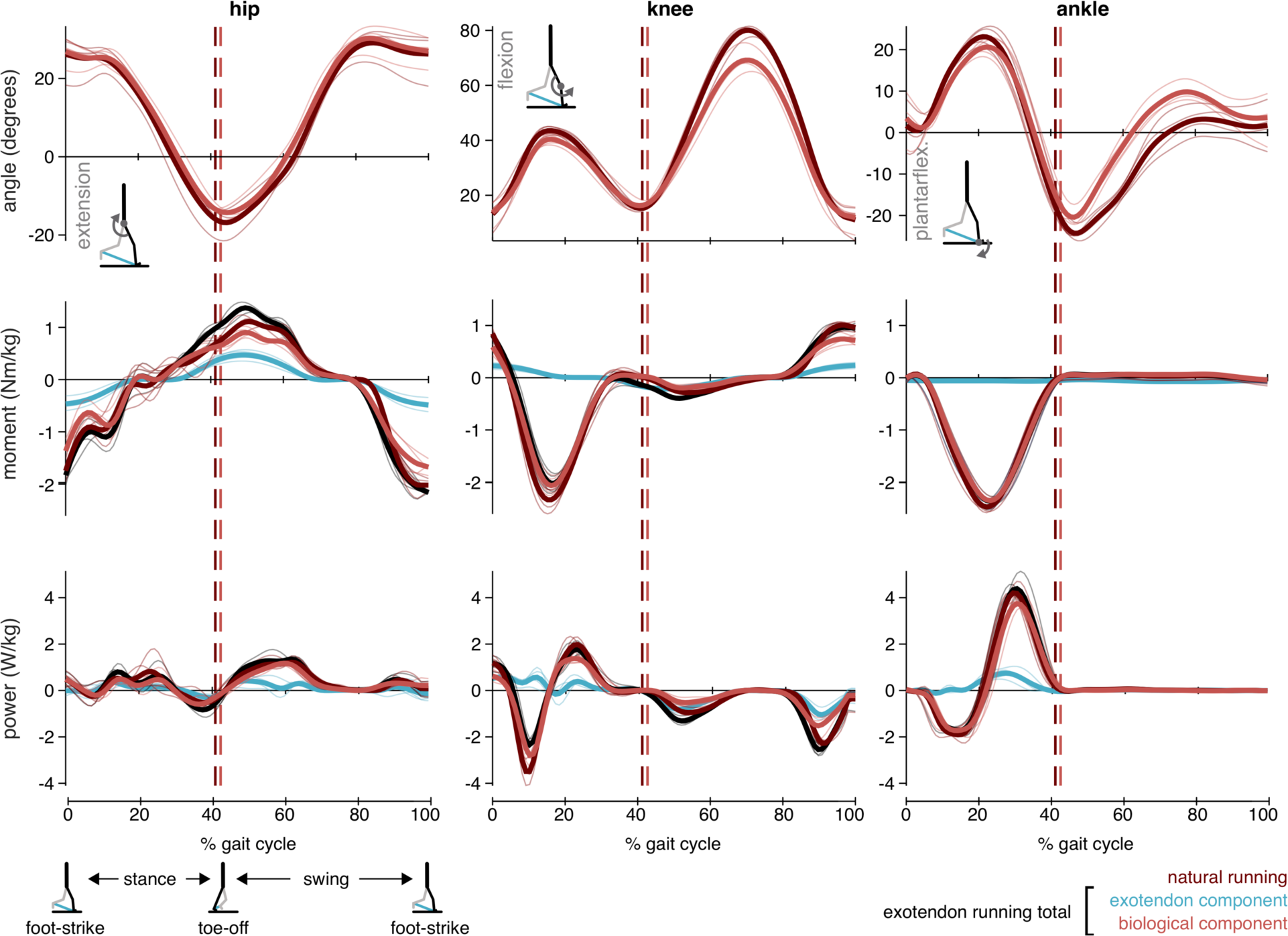
Joint-level kinematics and kinetics. Traces show average joint angles, moments, and powers across the gait cycle for natural running (dark red) and exotendon running. Kinetics from exotendon running are separated into exotendon contributions (blue), biological tissue (muscles, tendons, etc.) contributions (light red), and the total joint kinetics (black) for the four participants from Experiment 3. Thin traces show stride-averaged trajectories for individual participants (n=4) while the thick traces show trajectories averaged across participants. Vertical dashed lines indicate across-participant average toe-off time for exotendon running (light red) and natural running (dark red).

**Fig. S5.**
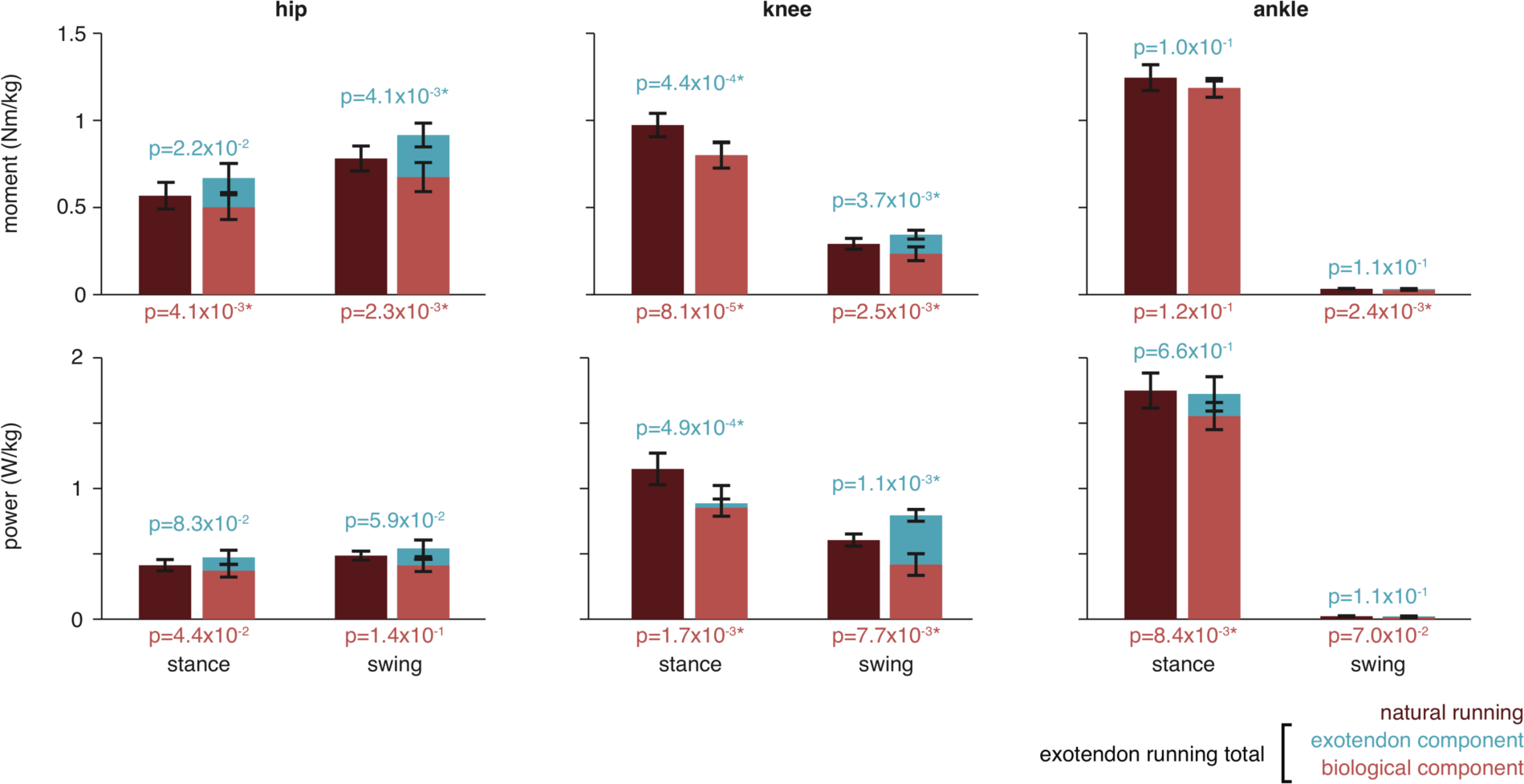
Average joint-level kinetics. Comparisons of average, absolute joint moments and powers across stance and swing for the four participants from Experiment 3. We compared moments and powers produced during natural running (dark red) to those produced during exotendon running. Average kinetics during exotendon runs were separated into the exotendon contribution (blue) and the biological contribution (light red). We report the p-values resulting from two-tailed paired t-tests comparing biological contributions to kinetics in natural and exotendon running below the axes (light red text) and comparing total kinetics in natural and exotendon runs above the bars (light blue). Asterisks indicate comparisons that were significant after Holm-Šidák corrections (alpha = 0.05). When running with the exotendon, during swing, hip, knee and ankle biological moments are reduced compared to natural running, as is knee power. During stance, hip and knee biological moments are reduced, along with knee and ankle powers. These reductions in biological moments suggest savings are achieved in both swing and stance. Total moments at the hip and knee, as well as total knee power, increased, demonstrating that adopting these kinematics without an exotendon would require additional effort compared to natural running.

**Fig. S6.**
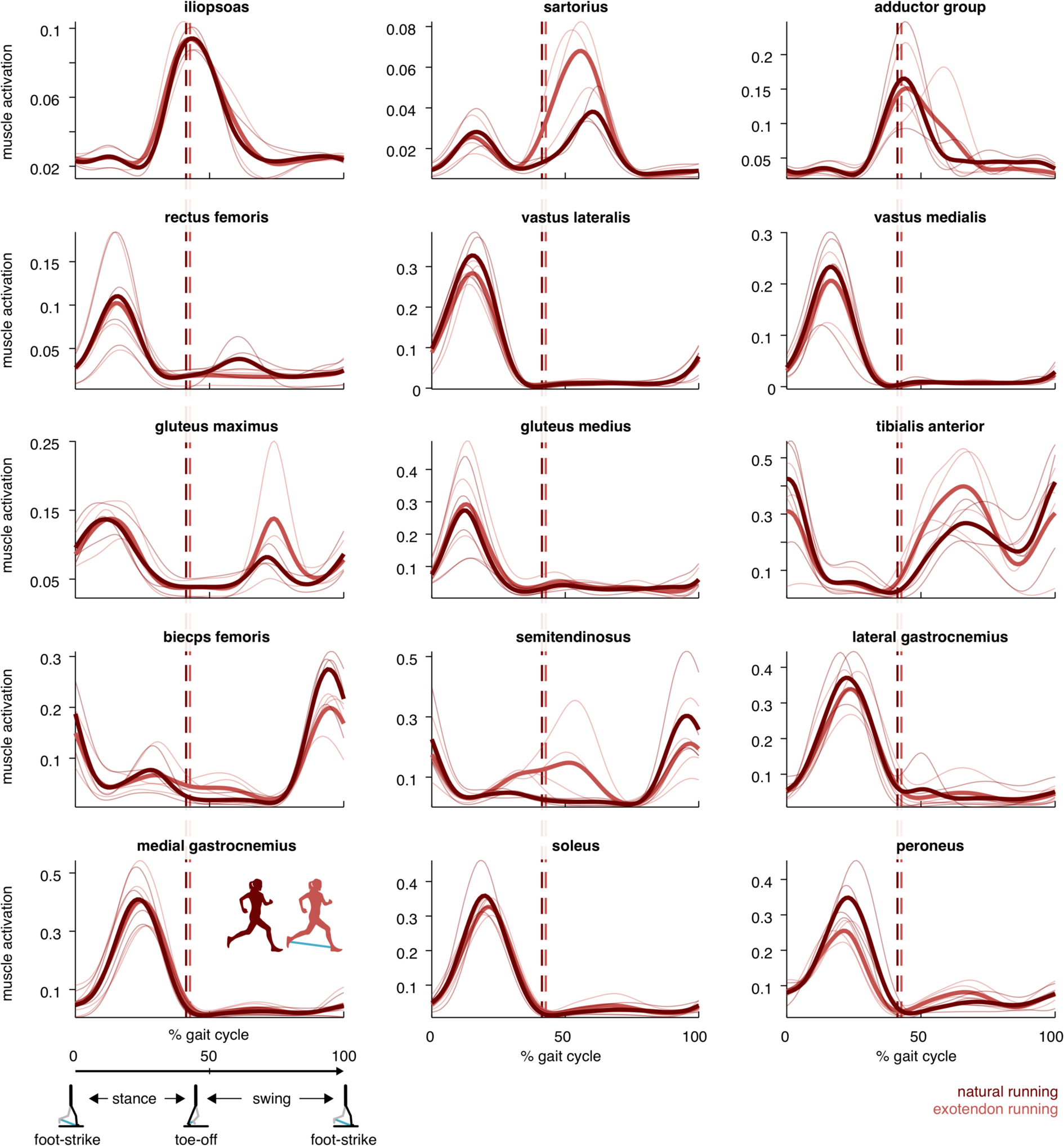
Muscle activity. Average muscle activity for each participant (thin traces) and across participants (solid lines) for natural running (dark red) and running with the exotendon (light red) as a function of gait cycle. The vertical dashed lines indicate the average time at which the toe lifts off the ground during exotendon running (light red) and natural running (dark red).

**Fig. S7.**
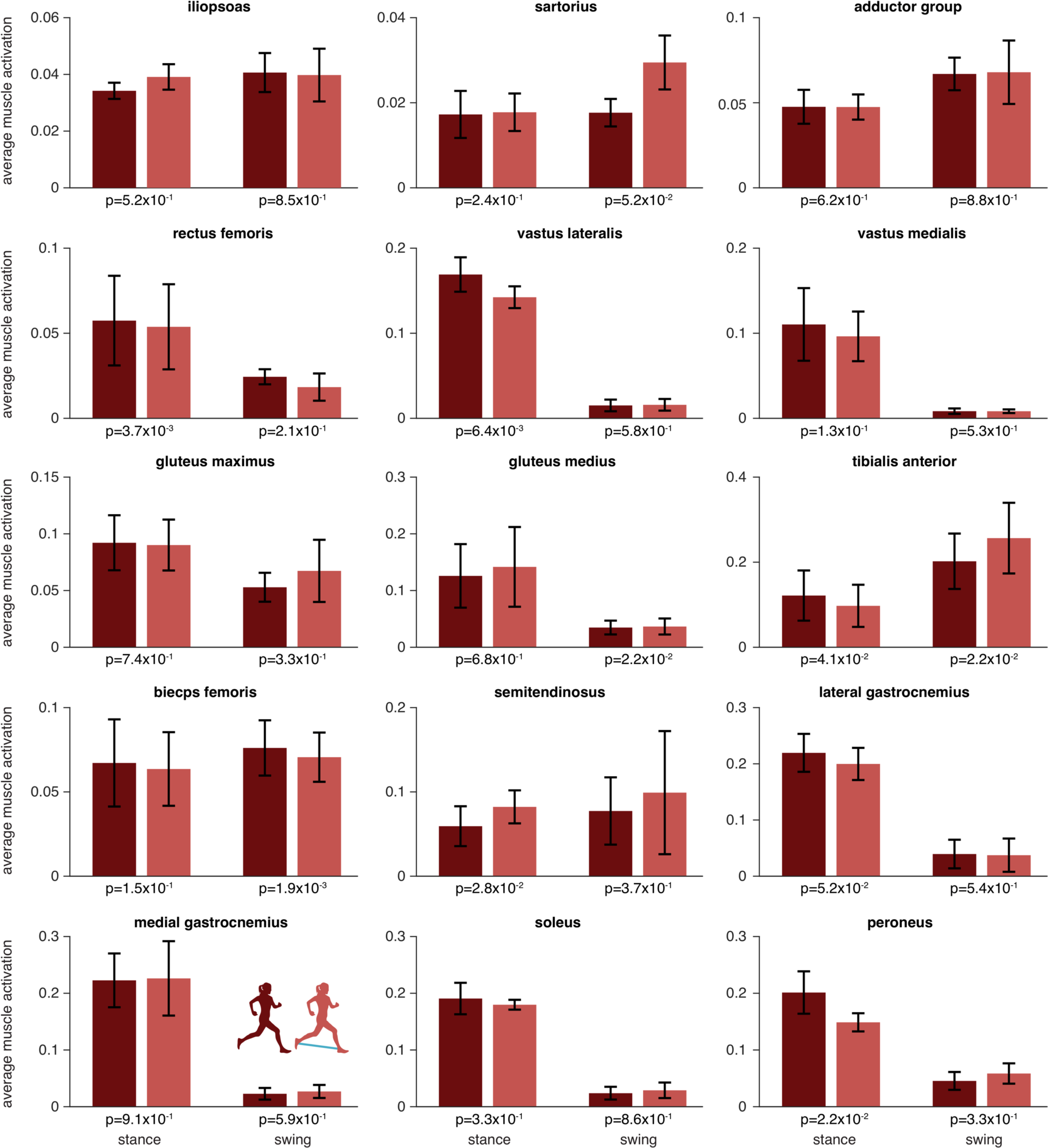
Average muscle activity. Comparisons of average, normalized muscle activity, computed from EMG recordings, across stance and swing phases of gait. Statistical comparison (paired t-test with Holm-Šidák corrections, α = 0.05) revealed no significant changes in muscle activity as a result of running with the exotendon.

**Fig. S8.**
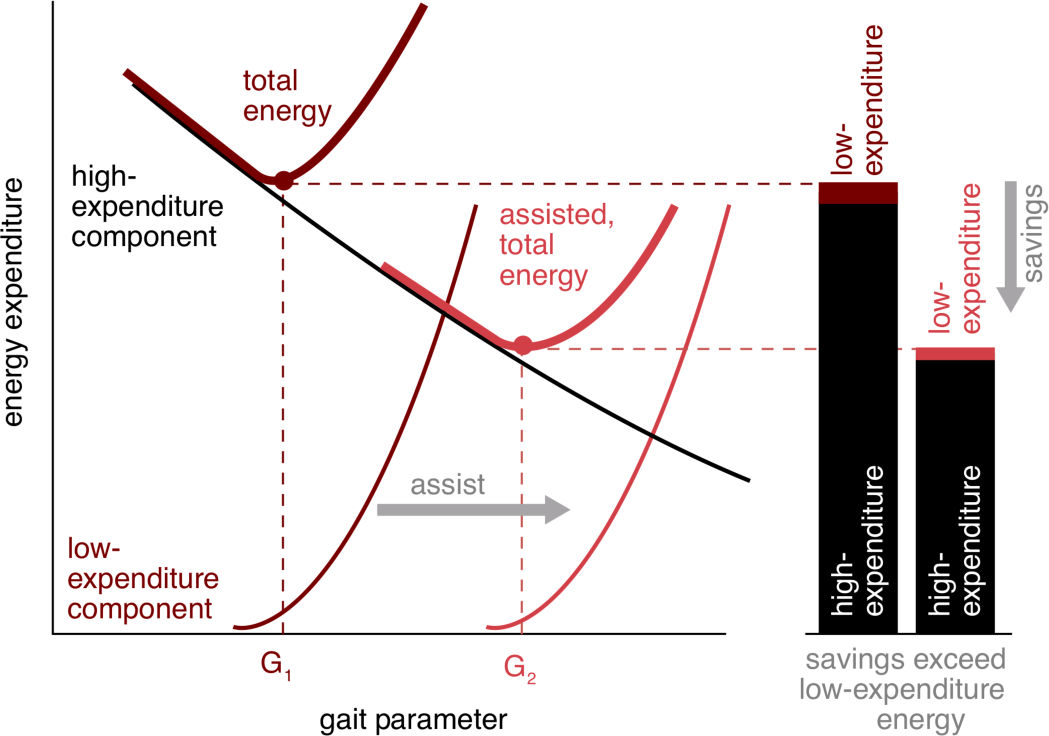
Targeting low-expenditure components of gait can result in savings that far exceed the expenditure associated with that component. Under natural running conditions, the optimal value of the gait parameter (G1) occurs at the minimum of the total energy expenditure curve (dark red bold), which is the sum of the low-expenditure component curve (dark red fine) and the high-expenditure component curve (black fine). Note that the low-expenditure curve has a steep slope beyond the optimal value of the gait parameter, preventing an otherwise favorable increase in the gait parameter. If an assistive device shifts the low-expenditure curve rightward, the optimal value of the gait parameter increases (G2). At this new optimum, the expenditure associated with the high-expenditure component is reduced by an amount that can greatly exceed the expenditure associated with the low-expenditure component.

